# The Impact of Memory and Stress on Choice Consistency

**DOI:** 10.1101/2024.12.09.627523

**Authors:** Fei Xin, Jialuo Lai, Manru Guo, Qingfei Chen, Jianhui Wu

**Affiliations:** School of Psychology, Shenzhen University, Shenzhen, China

**Author notes:** **Corresponding author:** Fei Xin, School of Psychology, Shenzhen University, Nanhai Ave 3688, Shenzhen 518060, Guangdong, China.

**Keywords:** choice consistency, memory, stress, fMRI, decision making

## Abstract

Choice consistency is a fundamental aspect of rational decision-making, reflecting the stability and reliability of an individual’s preferences. However, real-world decision-making often deviates from this ideal, as individuals frequently make irrational or inconsistent choices in value-based decision-making. This study combined computational modeling, neuroimaging, and behavioral assessments to elucidate the mechanisms by which stress and memory affect choice consistency. Remembered items exhibited higher choice consistency compared to forgotten items. Computational modeling further indicated that the drift rate was higher, and the decision threshold lower, for remembered food items compared to forgotten ones. Stress was found to impair both choice consistency and memory retrieval, with stress-induced declines in memory accuracy positively correlating with reductions in choice reaction times. Activation of the dorsolateral prefrontal cortex (DLPFC) during the pre-choice anticipation period was positively associated with choice consistency. Similarly, activation of the orbitofrontal cortex (OFC) during the memory retrieval of food stimuli correlated with improved memory accuracy. These findings suggest that stress may impair choice consistency by disrupting memory retrieval processes. Overall, our study provides novel insights into the role of stress and memory in decision-making, offering a more nuanced understanding of the neural and cognitive processes that govern choice behavior.

## 1. Introduction

Every day, individuals are confronted with numerous choices, such as deciding between eating cake or vegetables, and whether to drive or take the subway. In value-based decision-making, individuals evaluate available options based on their subjective values. By consistently selecting the option with the highest subjective value, rational individuals aim to achieve the greatest perceived benefit. According to rational choice theory, individuals make decisions to maximize their happiness or utility (Hodgson, 2012). Choice consistency is a fundamental aspect of rational decision-making, reflecting the stability and reliability of an individual’s preferences (Sugden, 1991).

However, real-world decision-making often deviates from this ideal, exhibiting variability that challenges traditional economic and psychological models, as individuals frequently make irrational or inconsistent choices (Kurtz-David et al., 2019; Simon, 1990). One potential source of this variability is the role of memory processes in shaping and maintaining choice consistency (Nitsch & Kalenscher, 2021; Wimmer & Shohamy, 2012). Episodic memory serves as a repository of personal experiences, providing a rich source of information that can be drawn upon when making decisions. Accurate and detailed memories may enable individuals to evaluate options based on past experiences, leading to more consistent and rational choices. The interplay between memory and choice consistency is particularly intriguing, as it suggests that the stability of preferences may depend on the accuracy and robustness of memory retrieval.

Recent studies have begun to explore the relationship between memory and value-based decision-making (Minxha et al., 2020; Michael N. Shadlen & D. Shohamy, 2016). However, only a limited number of studies have examined the influences of memory on choice consistency. Levin et al. (2019) investigated the relationship between age-related memory decline and inconsistencies in value-based decisions among 30 cognitively healthy older adults. They found that older adults with lower memory performance exhibited greater inconsistencies between their stated preferences and actual choices. In their study, memory performance was assessed using a cognitive assessment battery, rather than direct measurements of choice-relevant memory retrieval. Nitsch and Kalenscher (2021) employed a novel multi-attribute visual choice paradigm to experimentally test the influence of memory retrieval of exemplars on choice consistency. However, their results indicated a floor effect, suggesting low data quality for conclusively evaluating their hypotheses. Furthermore, their study, like others in the field, did not explore the neural mechanisms and temporal dynamics underlying the decision-making process. Therefore, more in-depth research is necessary to examine the association of memory with choice consistency.

Acute stress, characterized by a sudden and intense response to a perceived threat or challenge, can significantly alter cognitive processes, including memory retrieval and decision-making consistency. The impact of acute stress on episodic memory is well-documented, with research indicating that stress can lead to fragmented or distorted recollections (Gagnon et al., 2019; Shields et al., 2017). This phenomenon is largely attributed to the stress-induced release of glucocorticoids, which adversely affect the hippocampus, a brain region essential for memory formation and retrieval. Consequently, individuals under acute stress may struggle to accurately recall past events, which can have profound implications for their decision-making processes.

Research on the influence of acute stress on choice consistency remains sparse. The mechanisms underlying this effect are complex, involving the interplay of stress hormones, neural circuits, and cognitive strategies. Stress has been shown to trigger a shift from an analytic reasoning system to intuitive processes, potentially exacerbating behavioral biases in decision-making (Yu, 2016). Additionally, acute stress impairs self-control, cognitive flexibility, and episodic memory retrieval. These impairments raise critical questions: Does acute stress disrupt choice consistency? Does it do so by impairing the retrieval of option values from memory? Further investigation is needed to explore this potential relationship and to understand the underlying mechanisms that may contribute to decision-making inconsistencies under stress.

To investigate the aforementioned questions, 44 participants were recruited to perform memory and food choice tasks during fMRI scanning. Additionally, we applied a Hierarchical Drift Diffusion Model (HDDM) to the choice consistency and reaction times (RTs) obtained from all participants to assess how memory retrieval and acute stress affected choice consistency.

## 2. Materials and Methods

### 2.1. Participants

A priori power analysis using G∗Power (version 3.1) indicated that a sample size of 36 would allow for the detection of medium effect size (*f* = 0.25) with 95% power at an alpha of 0.05 for the repeated measures with two within-subject factors (Stress: No-Shock vs. Shock; Memory: remembered vs. forgotten). *N* = 44 healthy, right-handed participants were enrolled in the study (age range: 18 to 27; mean age = 20.64 ± 2.33 years; 22 females). One participant was excluded from the HDDM analysis due to a lack of Forgotten trials in the No-Shock condition. All participants were free from current or past psychiatric, neurological, or other medical disorders (self-reported). Participants were instructed to abstain from alcohol and caffeine for 24 hours prior to the experiment. Written informed consent was obtained after a detailed explanation of the study protocol. The study procedures received full ethical approval from the local ethics committee and were conducted in accordance with the latest revision of the Declaration of Helsinki.

### 2.2. Experimental design and procedure

The experiment consisted of four phases: (1) rating phase 1, (2) rating phase 2, (3) memory distraction, and (4) the memory and choice task. Hunger ratings were collected at two times during the study: (1) before the food rating task, and (2) before the memory and choice task. Participants rated their overall hunger (i.e., How hungry do you feel?) on a 9-point Likert scale from 1 (not at all) to 9 (very much). Figure 1 provides a schematic overview of the experimental protocol.

**Figure 1.**
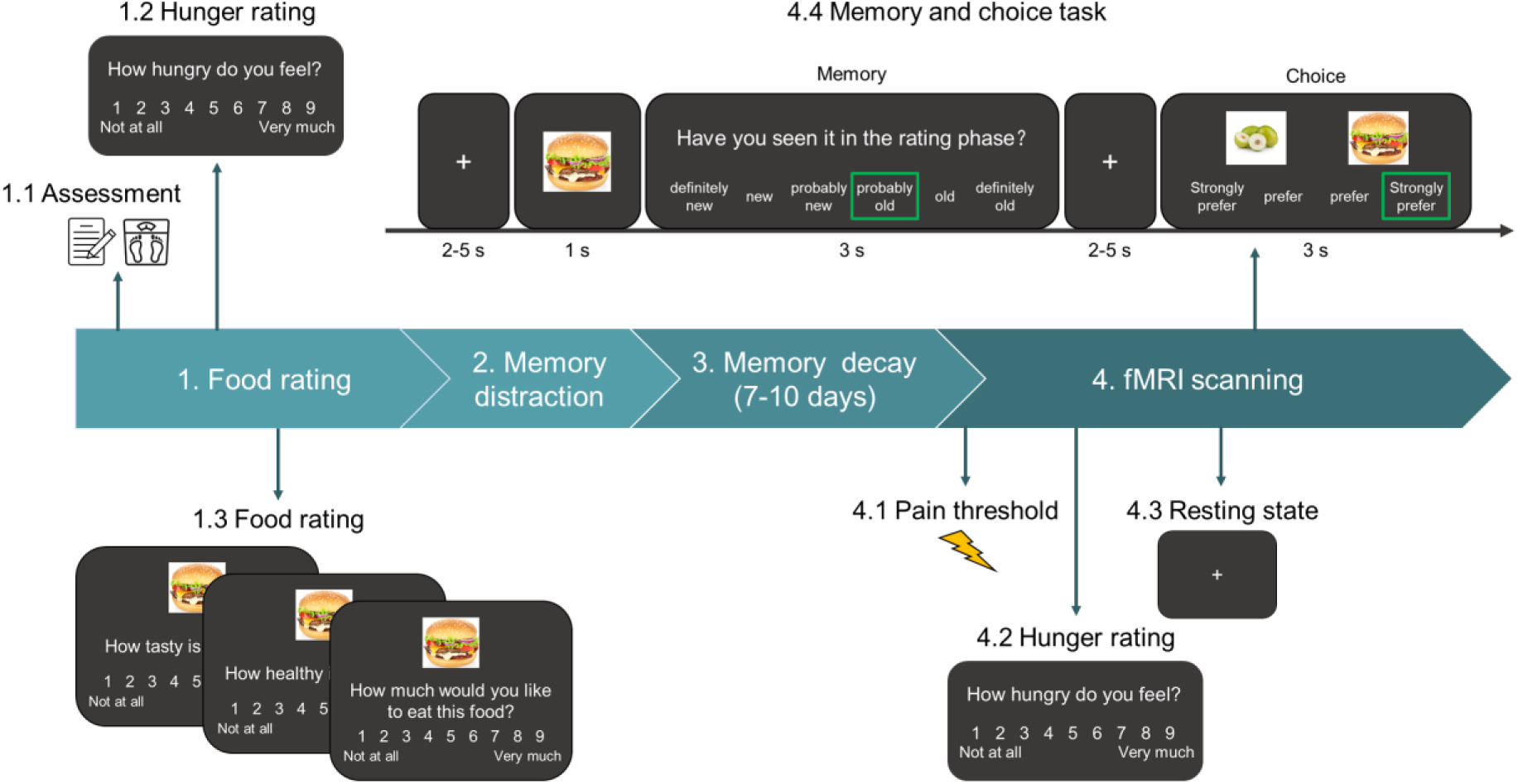
Study design and experimental task. The scheme illustrates the temporal organization of the experiment with the different steps indicated by the numbers 1-4. The duration of the fMRI task is shown in seconds.

#### Rating phase 1

Participants were presented with 186 food items and asked to rate each item on a generalized visual analog scale, ranged from 1 (not at all) to 9 (very much). Over three separate runs, participants rated the foods in terms of taste (“How tasty is this food?”), health (“How healthy is this food?”), and craving (“How much would you like to eat this food at the end of the experiment?”). The order of the taste and health runs was counterbalanced among the participants, while the craving run always came last. The order of stimulus presentation was randomized in each task. All food items were familiar to the participants, and easily available from nearby supermarkets. Participants had viewed all the food images before the ratings to effectively use the full range of the rating scale (Brus et al., 2021).

#### Rating phase 2

To ensure the reliability of participants’ value ratings and to identify excessive variability, rating phase 2, identical to rating phase 1, was conducted immediately after phase 1. Participants were not informed in advance about rating phase 2, a deliberate measure to prevent them from consciously memorizing or attempting to replicate their slider positions from phase 1. Participants whose ratings displayed significant inconsistencies across the two phases were excluded from further analysis.

#### Memory distraction

To prevent rehearsal of information obtained during the encoding phase, a memory distraction stage was conducted immediately after two rating phases. Participants first played an online game called “Spot the Difference” (https://www.spotthedifference.com/trocadero), followed by a facial 2-back task (Xin & Lei, 2015).

#### Decision-making task

The MRI acquisition was carried out 7-10 days after rating and memory distraction phases, including an 8 min 20 s resting-state scan (Rest), and six decision-making task runs. During the resting-state fMRI acquisition, participants were instructed to passively view a fixation cross and keep as motionless as possible. Brain structural data was acquired after three decision-making task runs. Head movements were minimized by using comfortable head cushions.

Each decision-making trial includes a memory phase and a choice phase. Trials began with a jittered fixation cross for 2–5 s, followed by the presentation of one food item for 1 s and a 3 s recognition memory phase. Participants were asked to indicate whether they had seen the food item during the previous rating phases on a 6-point confidence scale (1 = “definitely new”, 2 = “new”, 3 = “probably new”, 4 = “probably old”, 5 = “old”, 6 = “definitely old”). A total of 180 food items (108 old, 72 new) were randomly mixed together. After the memory phase, a jittered period of 2–5 s preceded the choice phase. During the choice phase, participants were presented with the reference food and a trial-unique food on 180 trials. For each participant, a reference food item rated as median (i.e., 5) for craving was selected at random. Another 180 food items were also selected, half with higher craving scores and half with lower craving scores than the reference food. The reference food was always presented on the same side of the screen (left or right), with the position balanced between participants. Participants indicated their preference for one of the two items using a Likert scale with “strongly prefer” anchoring each end. To incentivize participants to make choices according to their real preferences, they were told they would receive a snack-sized portion of one of their chosen foods, selected at random, after the task.

To investigate the potential influence of a concurrent stressor on the neural and cognitive mechanisms underlying choice consistency, participants completed the decision-making task under two conditions: threat of shock (three runs) and safety (No-Shock; three runs). At the beginning of each task run, participants were informed whether the condition was “unpredictable Shock” or “No-Shock”. Fixations in the shock run were red, while those in the No-Shock run were white. At the end of each run, participants rated the intensity of their emotional experience (e.g., “How anxious/fearful were you during threat/safe?”) and their level of attentional control (e.g., “How difficult was it to pay attention to the task during threat/safe?”) using 9-point scales (”not at all” to “extremely”).

### 2.3. Stress Manipulation

Acute stress was induced using a well-established threat-of-shock procedure (Schmitz & Grillon, 2012). Shocks were delivered for 50 ms to the participants’ left ankle using SXC-4S multichannel electrical stimulator via two electrodes. Participants rated the level of pain evoked by the shocks on an 11-point scale (0 = no sensation, 10 = maximum tolerable pain). Each participant underwent an individual titration procedure to calibrate the level of shock. Pain calibration began with a weak electric shock (0.5 mA) and was gradually adjusted until a rating of 8 was reported. This intensity was used throughout the decision-making task. Participants were instructed that this selected level of shock would be delivered for brief durations during uncertain periods of the task, and that the timing would be unrelated to their performance on the task. During each “threat” run, participants were explicitly told they would receive between 6 and 12 shocks, ensuring a sense of unpredictability. To prevent the shocks from interfering with task performance, all shocks were administered during the fixation period.

### 2.4. Psychometric assessment

Beck’s Depression Inventory (BDI-II) (Beck et al., 1996) was administered to evaluate participants’ level of depression. Before and after MRI scanning, participants completed the State-Trait Anxiety Inventory (STAI) (Spielberg et al., 1983) and the Positive and Negative Affect Schedule (PANAS) (Watson et al., 1988) to evaluate their current emotional state. All participants were in the normal range with respect to depression and anxiety scores confirming the self-reported lack of psychiatric conditions in the present sample (Baron-Cohen et al., 2001; Beck et al., 1996).

### 2.5. Behavioral analyses

Food items recognized with high confidence (scored 5 or 6) were classified as remembered items (R), while those scored 4 or lower were classified as forgotten items (F). Items scored 4 (i.e., participants guessed they were “probably old” items) were considered as forgotten items to balance the numbers of remembered and forgotten trials and to maximize statistical power (Zheng et al., 2018). A consistent choice was defined as a trial in which the participant chose the item they had assigned a higher craving rating (Brus et al., 2021). Trials with reaction times below 300 ms were discarded as outliers.

### 2.6. Computational modeling

Hierarchical Drift Diffusion Model (HDDM) analysis was conducted to decompose behavioral responses into underlying cognitive processes. The decision-making process is modeled as the gradual accumulation of information, making a threshold decision when the accumulated information surpasses a predefined threshold. The model typically returns four parameters: drift rate (v, representing information acquisition speed), threshold (α, representing the distance between two selection boundaries), starting point (z, representing the prior deviation of a response), and nondecision time (τ, representing the duration of the nondecision process).

We fitted the participants’ food decision-making performance (i.e., accuracy and reaction time) with an HDDM (Wiecki et al., 2013), which assumes a stochastic accumulation of sensory evidence over time, toward one of two decision boundaries corresponding to consistent and inconsistent choices. Specifically, we defined a consistent response as the upper boundary and an inconsistent response as the lower boundary. A larger threshold between the two boundaries indicates a more conservative decision-making strategy, requiring more evidence to make a decision. A higher absolute value of drift rate indicates faster information accumulation, leading to quicker decision-making. The starting point represents the initial position where evidence begins to accumulate; a starting point greater than α/2 indicates a bias towards one decision boundary. Nondecision time accounts for processes other than information accumulation, such as information encoding and action execution.

We implemented HDDM using the HDDM v1.0.1 package (Wiecki et al., 2013) in Python v3.9. Three parameters were chosen for fitting: drift rate (v), threshold (α), and nondecision time (τ), with nondecision time varying among participants. Additionally, we established 15 models for drift rate and threshold based on the main effects and interactions of two variables: stress and memory. Model comparison was conducted using Deviance Information Criterion (DIC), Widely Applicable Information Criterion (WAIC), and Pareto-Smoothed Importance Sampling Leave-One-Out Cross-Validation (LOOCV) to select the best-fitting model (Pan et al., 2022).

### 2.7. fMRI data acquisition

MRI data was collected on a 3T Siemens Prisma scanner (Siemens Healthineers, Erlangen, Germany). High-resolution brain structural data was acquired using a T1-weighted sequence (repetition time [TR] = 2530 ms, echo time [TE] = 2.98 ms, flip angle = 7°, thickness = 1 mm with no gap, field of view [FOV] = 256 mm × 256 mm, acquisition matrix = 224 × 224, reconstruction matrix = 448 × 448, in-plane resolution = 0.5 × 0.5 mm^2^, thickness = 1 mm, slices = 192) to improve spatial normalization of the functional data. Functional data were acquired using a T2*-weighted gradient Echo Planar Imaging (EPI) sequence with the following parameters: TR/TE = 2000/30ms, flip angle = 90°, acquisition matrix = 112 × 112, FOV = 224 × 224 mm^2^, thickness/gap = 2/0.3 mm, in-plane resolution = 2.0 × 2.0 mm^2^, axial slices = 62, number of multiband acceleration factor = 2.

### 2.8. fMRI data analysis

fMRI images were preprocessed and analyzed using the standard procedure in SPM12 (http://www.fil.ion.ucl.ac.uk/spm/; Wellcome Trust Centre for Neuroimaging) and DPABI (Version: V5.4_230110; http://rfmri.org/dpabi)(Yan et al., 2016). The first 5 volumes of each functional run were removed to allow for MRI signal equilibrium. The preprocessing steps included slice timing, head motion correction, spatial normalization and smoothing (6 mm full-width at half maximum [FWHM] Gaussian kernel). For each participant and each run, a general linear model (GLM) was constructed by convolving the regressors with the canonical hemodynamic response function (HRF) to estimate evoked blood-oxygen level-dependent (BOLD) activity. The GLM contained regressors indicating the food recognition (duration = 1 s), recognition rating (duration = 3 s), anticipatory choice (duration = 2-5 s, subject-specific time), choice (duration = 3 s), and six motion-correction parameters. All the results were corrected for multiple comparisons using family-wise error (FWE) rate at *p < 0.05* voxel-level (vFWE) or cluster-level (cFWE, cluster-forming threshold *p* < 0.001).

To investigate the associations between the BOLD activity and the behavioral performance, parameter estimates were extracted from a 4-mm radius sphere centered at the peak coordinate of the regions of interest (ROIs). ROIs included the orbitofrontal cortex (OFC; MNI = [−20, 44, −16]; Table S7) for the contrast remembered versus forgotten food stimuli, and the dorsolateral prefrontal cortex (DLPFC; MNI = [−38, 36, 28]; Table S5) for the contrast consistent versus inconsistent anticipatory choices. The OFC was selected based on previous studies demonstrating its involvement in memory for food stimuli (Morris & Dolan, 2001). The DLPFC was chosen based on studies showing its role in value-based decision making and choice consistency (Hutcherson & Tusche, 2022; Zyuzin et al., 2023). Pearson correlations were calculated between the mean parameter estimates from the ROI clusters and behavioral performances. Additionally, time courses were extracted from the OFC to compare neural dynamics between remembered and the forgotten conditions, and from the DLPFC to compare neural dynamics between consistent and inconsistent conditions.

## 3. Results

### 3.1. Threat of shock induces aversive affect

Paired *t*-tests demonstrated that: 1)self-reported anxiety was higher (*t*[43] = 8.551, *p* < 0.001) during the Shock condition (mean ± SD = 4.758 ± 2.006) than the No−Shock condition (2.387 ± 1.360); 2) self-reported fear was higher (*t*[43] = 9.892, *p* < 0.001) during the Shock condition (4.977 ± 1.963) than the No-Shock condition (1.849 ± 1.185); 3) self-reported difficulty in concentrating was higher (*t*[43] = 5.204, *p* < 0.001) during the Shock condition (4.190 ± 1.546) than the No−Shock condition (2.856 ± 1.584).

### 3.2. Influences of stress and memory on choice consistency

The interaction effect of memory and stress on choice consistency were assessed using a two-way repeated measures analysis of variance (ANOVA) with two within-subject factors: Memory (Remembered, Forgotten) and Stress (No-Shock, Shock). No significant Memory × Stress interaction effect was found for choice consistency (*p* = 0.772). However, significant main effects of Memory (*F*[1, 43] = 6.988, *p* = 0.011, *η^2^_P_* = 0.140) and Stress (*F*[1, 43] = 7.041, *p* = 0.011, *η^2^_P_* = 0.141) were observed (Figure 2A). For choice reaction time, no significant Memory × Stress interaction effect was observed (*p* = 0.165). Significant main effects of Memory (*F*[1, 43] = 24.233, *p* < 0.001, *η^2^_P_* = 0.360) and Stress (*F*[1, 43] = 22.628, *p* < 0.001, *η^2^_P_* = 0.345) were found (Figure 2B). Additionally, we investigated the impact of stress on memory performance and found that stress significantly reduced memory accuracy (*p* = 0.007; Figure 2C). The decline in memory accuracy due to stress was positively correlated with a reduction in reaction time for choice tasks (*r*[42] = 0.329, *p* = 0.029; Figure 2D). Reaction times for inconsistent choices were significantly longer than those for consistent choices (*t*[43] = 4.745, *p* < 0.001; Figure 2E).

**Figure 2.**
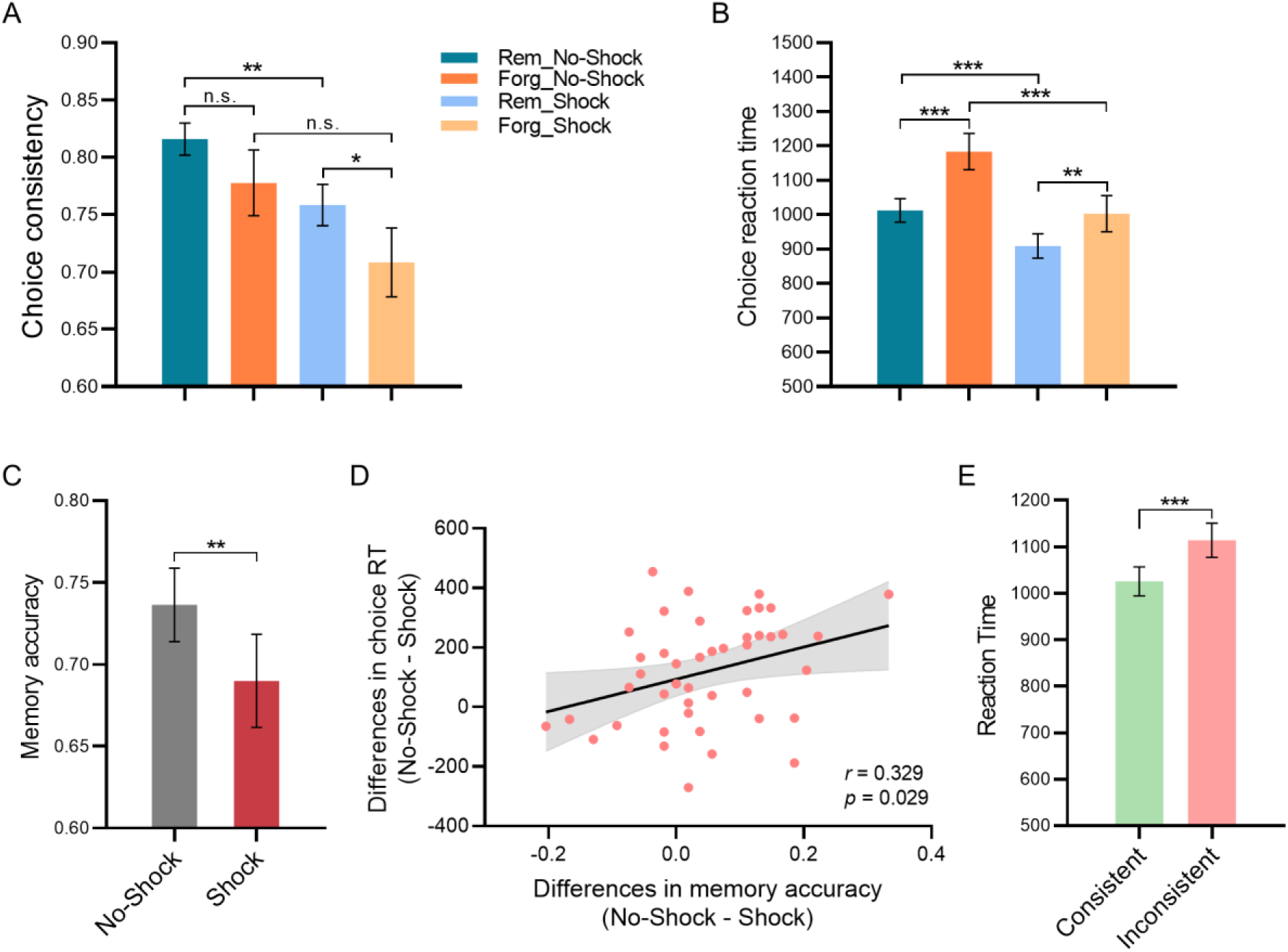
Behavioral performance (mean ± standard error) on the food choice task. A) Choice consistency for remembered and forgotten items under No-Shock and Shock conditions; B) Choice reaction times for remembered and forgotten items under No-Shock and Shock conditions; C) Memory accuracy under No-Shock and Shock conditions; D) The reduction in memory accuracy due to stress was positively correlated with the decrease in choice reaction time; E) Reaction times for consistent and inconsistent choices. n.s., not significant; \**p* < 0.05; \*\**p* < 0.01; \*\*\**p* < 0.001. Rem, remembered; Forg, forgotten.

### 3.3. Smaller value difference leads to greater choice inconsistency

Participants’ craving rating for the reference item were subtracted from craving rating for the target item to calculate a signed value difference (VD) between each pair of items. The decision trials were divided into four VD levels on the rating scale (−4 and −3, −2 and −1, 1 and 2, 3 and 4), as defined by the first rating provided by each participant (Figure 3A). Consistent with previous research (Alós-Ferrer & Garagnani, 2021; Brus et al., 2021; Kurtz-David et al., 2019), we found that smaller the value difference, the higher choice difficulty level. Difficult choices lead to greater choice inconsistency (Figure 3B) and longer reaction times (Figure 3C).

**Figure 3.**
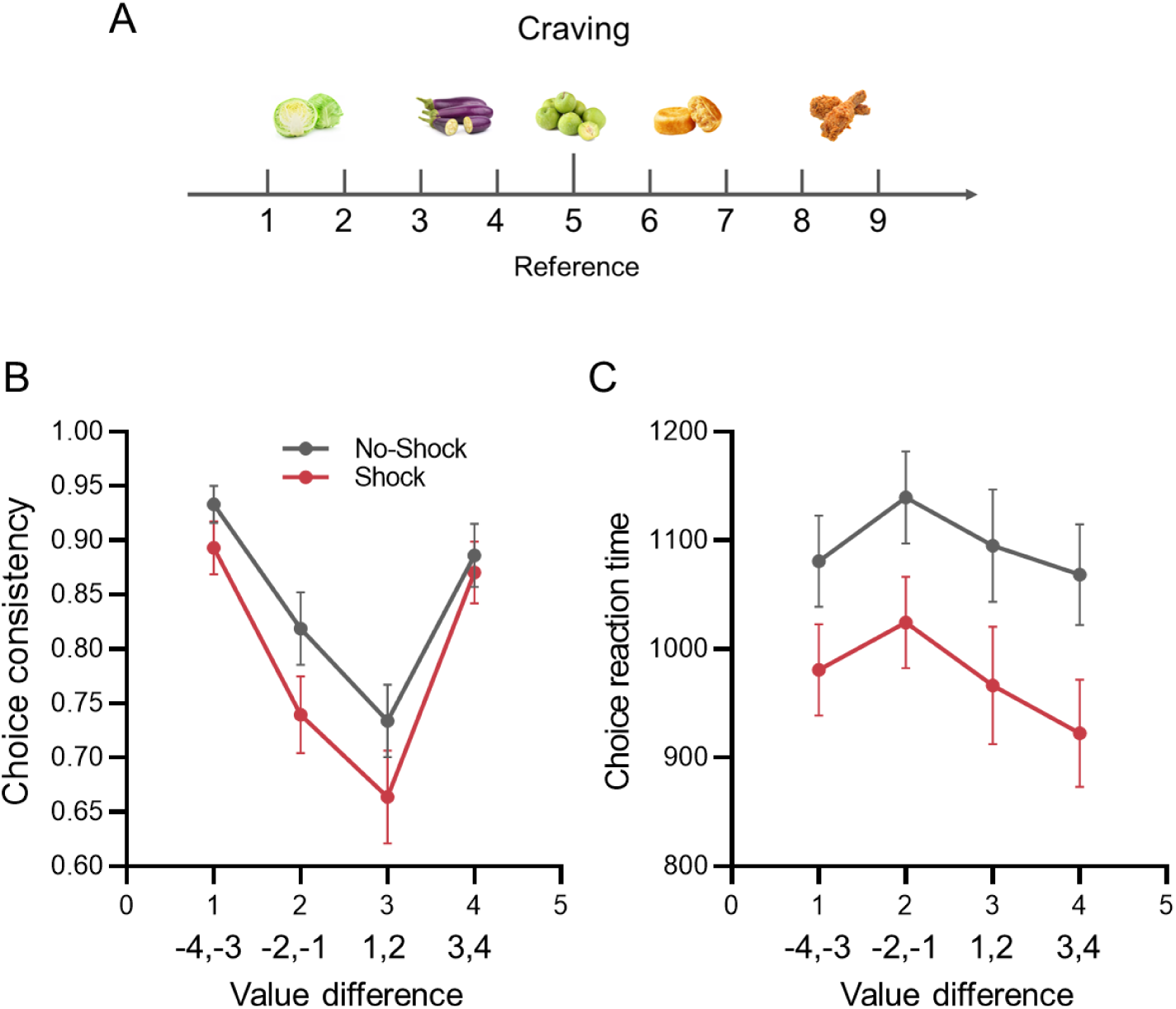
A) Distribution of craving ratings. B) Choices were more inconsistent when the value difference between alternatives was smaller. C) Participants made choices more slowly when the value difference between alternatives was smaller.

### 3.4. HDDM results

The best-fitting model was Model One, which achieved the optimal complexity-approximation trade-off, as evaluated by the DIC, WAIC, and LOOCV, with lower values indicating a better fit (see Table S1). In this model, the drift rate was influenced solely by the main effect of memory, while the decision threshold was affected by the main effects of both stress and memory, without interaction. By comparing the 95% Highest density interval (HDI) of the posterior difference distribution under different conditions at the group level, we can determine significant differences between conditions by checking if the interval does not include zero (Johnson et al., 2017).

The HDDM results indicates that, compared to Forgotten food items, the drift rate is higher for Remembered food items (*Mean* = 0.13, 95% HDI = [0.04, 0.21], excluding 0), and the threshold is lower (*Mean* = −0.12, 95% HDI = [−0.19, −0.06], excluding 0). Additionally, compared to the No-Shock condition, the threshold is lower in the Shock condition (*Mean* = −0.19, 95% HDI = [−0.28, −0.11], excluding 0). For more details, please refer to Figure 4.

**Figure 4.**
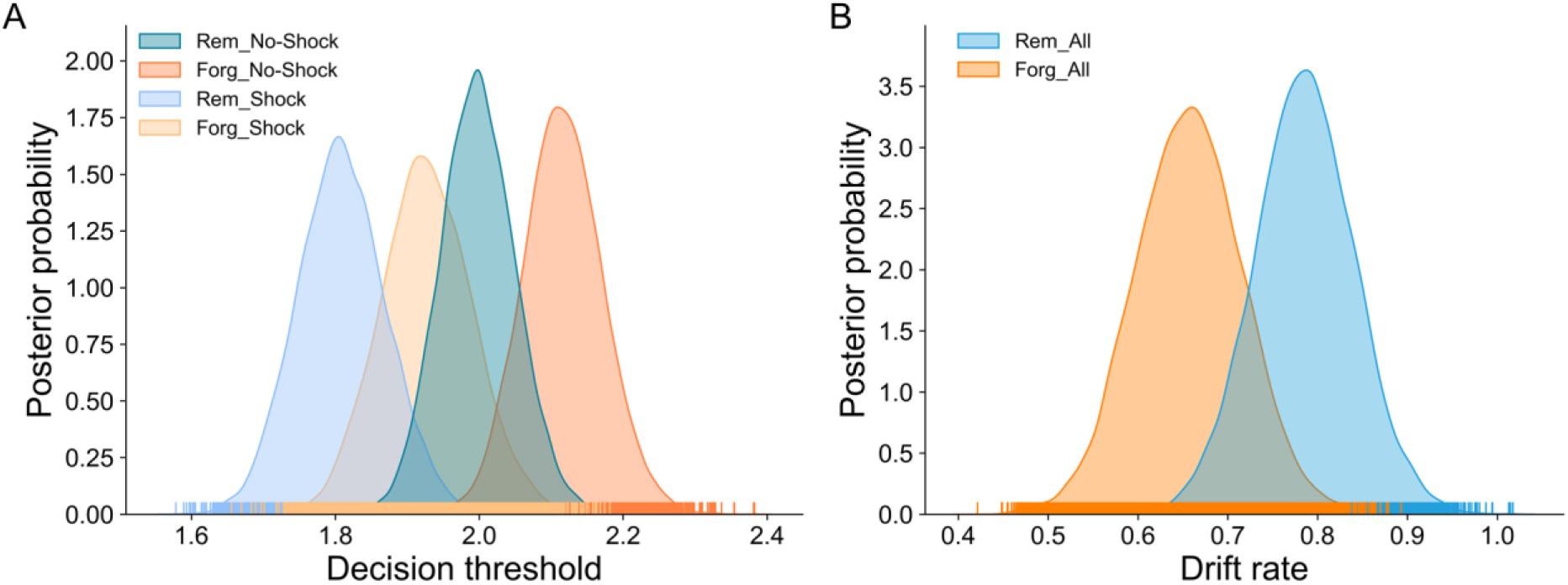
A) Group-level posterior probability distributions of threshold under different conditions. B) Group-level posterior probability distributions of drift rates under different memory conditions. Rem, remembered; Forg, forgotten.

### 3.5. fMRI results

#### 3.5.1. Neural correlates of choice consistency

The contrast of Consistent > Inconsistent revealed no significant brain regions during the choice period. To further elucidate the differences in brain activity between consistent and inconsistent choices, we conducted exploratory analyses. Time courses were extracted from the hippocampus, parahippocampal gyrus, OFC, and DLPFC during the choice period. The results indicated that activity in these four regions was significantly higher during consistent choices compared to inconsistent choices (Figure 5). The regions of interest (ROIs) for the hippocampus, parahippocampal gyrus, OFC, and DLPFC were created using the Automated Anatomic Labelling (AAL) atlas (Tzourio-Mazoyer et al., 2002), as implemented in WFU Pickatlas (Maldjian et al., 2003). Conversely, the contrast of Inconsistent > Consistent revealed activation in several distributed brain regions during the choice period, including the anterior cingulate cortex, supplementary motor area, inferior parietal lobule, middle frontal gyrus, superior frontal gyrus, inferior frontal gyrus, superior parietal lobule etc. (Figure S2, Table S1).

**Figure 5.**
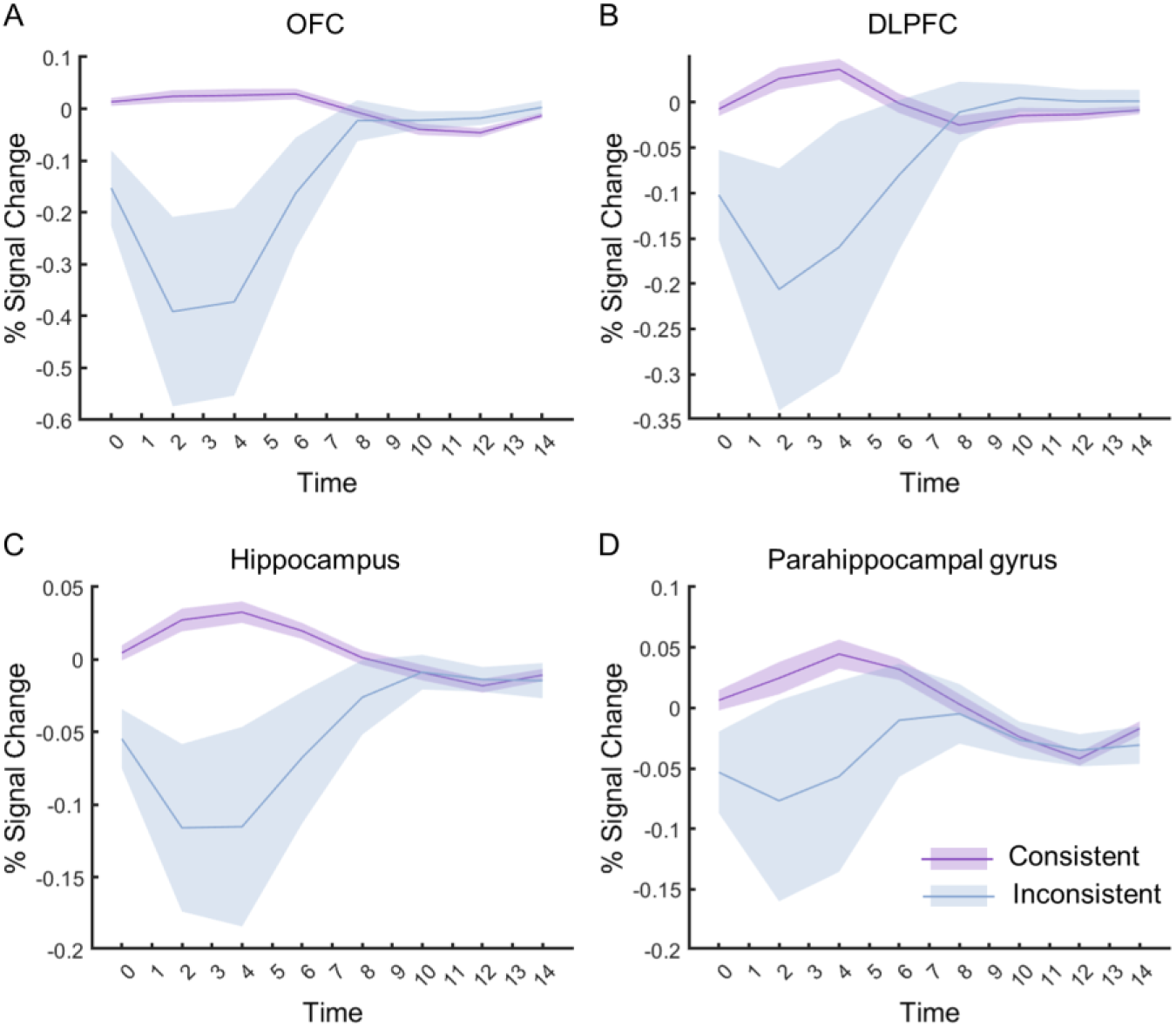
Time courses were extracted from atlas-based regions in the A) OFC, B) DLPFC, C) hippocampus, and D) parahippocampal gyrus.

#### 3.5.2. The effects of stress on choice consistency

To determine the impact of stress on choice consistency, we compared brain activation between stress (Shock) and non-stress (No-Shock) conditions during the choice period. The contrast of Shock > No-Shock revealed distributed brain regions with stronger activation under stress conditions, including the postcentral gyrus, precuneus, superior temporal gyrus, supramarginal gyrus, insula, lingual gyrus etc. (Figure 6A, Table S3). Conversely, the contrast of No-Shock > Shock showed brain regions with stronger activation under non-stress conditions, including the postcentral gyrus, precentral gyrus, fusiform gyrus, parahippocampa gyrus, hippocampus etc. (Figure 6B, Table S4).

**Figure 6.**
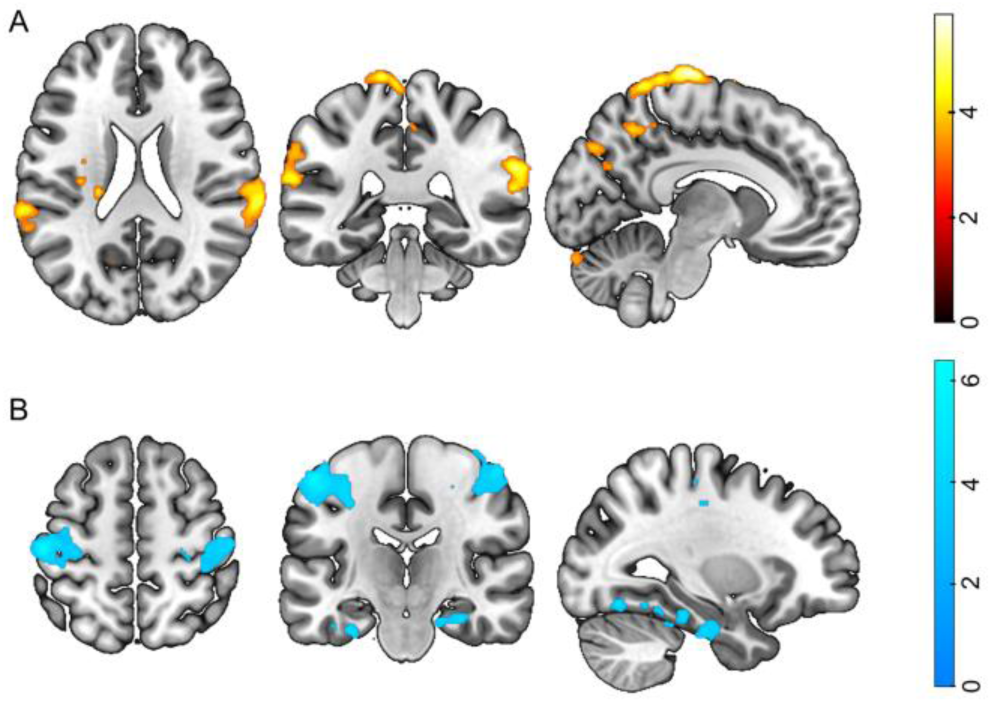
Whole-brain cluster-level family-wise error corrected group-level results (*p < 0.05*) for the contrasts A) Shock > No-Shock and B) No-Shock > Shock during the choice period.

#### 3.5.3. Neural responses during the anticipation of choice

The contrast of Consistent > Inconsistent across both No-Shock and Shock conditions revealed distributed brain regions with stronger activation for consistent choices compared to inconsistent choices during the pre-choice anticipation period. These regions included the DLPFC, middle cingulate, supramarginal gyrus, and inferior parietal lobule etc. (Figure 7A, Table S5). A significant positive association was observed between DLPFC activity with choice consistency (*r*[42] = 0.319, *p* = 0.035; Figure 7B). Time course extracted from the DLPFC indicated that consistent choices induced higher DLPFC activity than inconsistent choices during the pre-choice anticipation period (Figure 7C).

**Figure 7.**
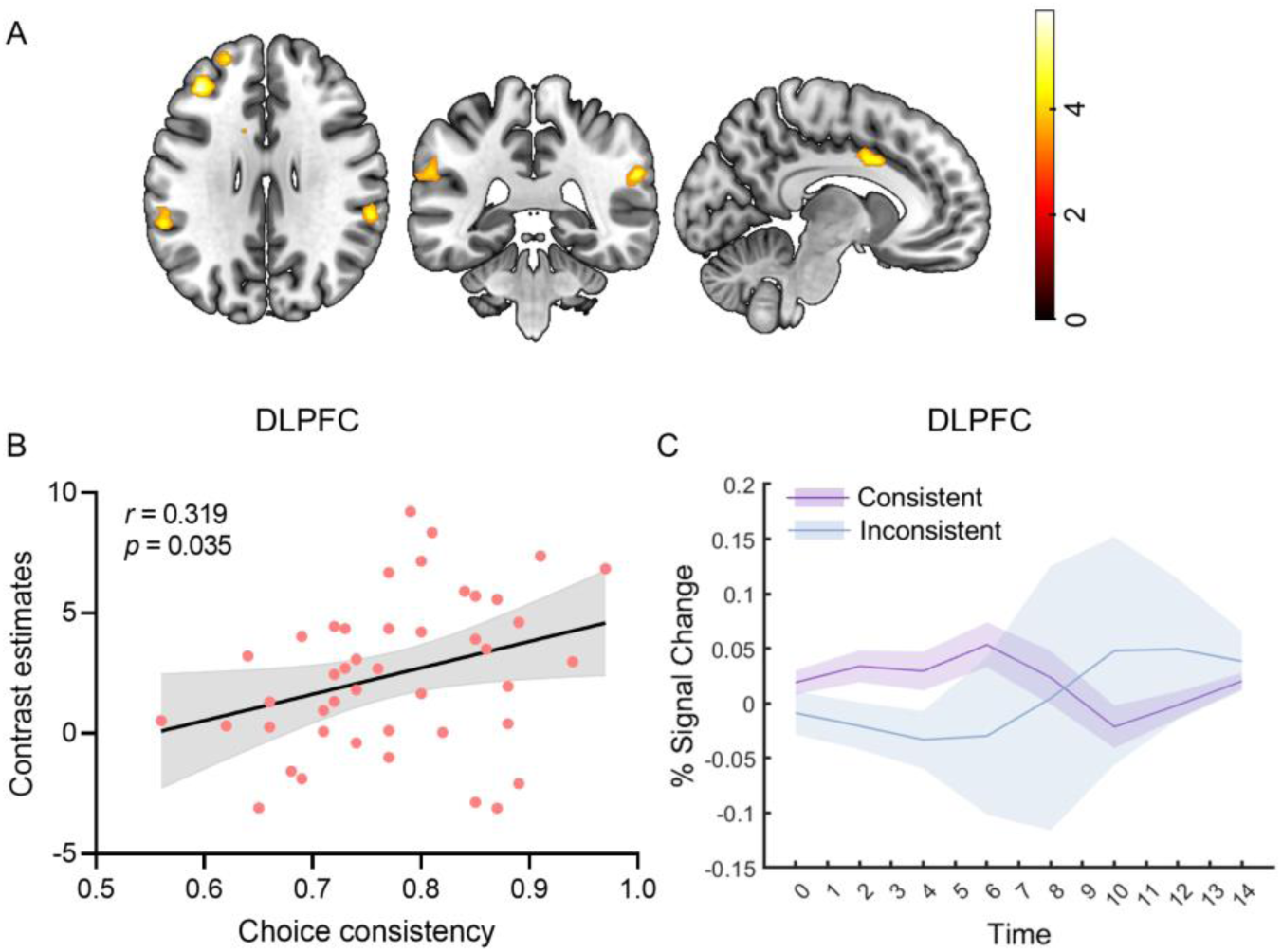
A) Whole-brain cluster-level family-wise error corrected group-level results (*p < 0.05*) for the contrast Consistent > Inconsistent during the pre-choice anticipation period; B) A significant positive correlation between DLPFC activity and choice consistency; C) Time course of the DLPFC in the Consistent and Inconsistent conditions during the pre-choice anticipation period.

#### 3.5.4. The effects of stress on anticipation of choice

To determine the impact of stress on the anticipation of choice, we compared brain activation between stress (Shock) and non-stress (No-Shock) conditions. The contrast of Shock > No-Shock revealed distributed brain regions with stronger activation under stress conditions, including the paracentral lobule, postcentral gyrus, middle cingulate, superior temporal gyrus, and insula etc. (Figure 8, Table S6). The contrast of No-Shock > Shock revealed no significant brain regions during the pre-choice anticipation period.

**Figure 8.**
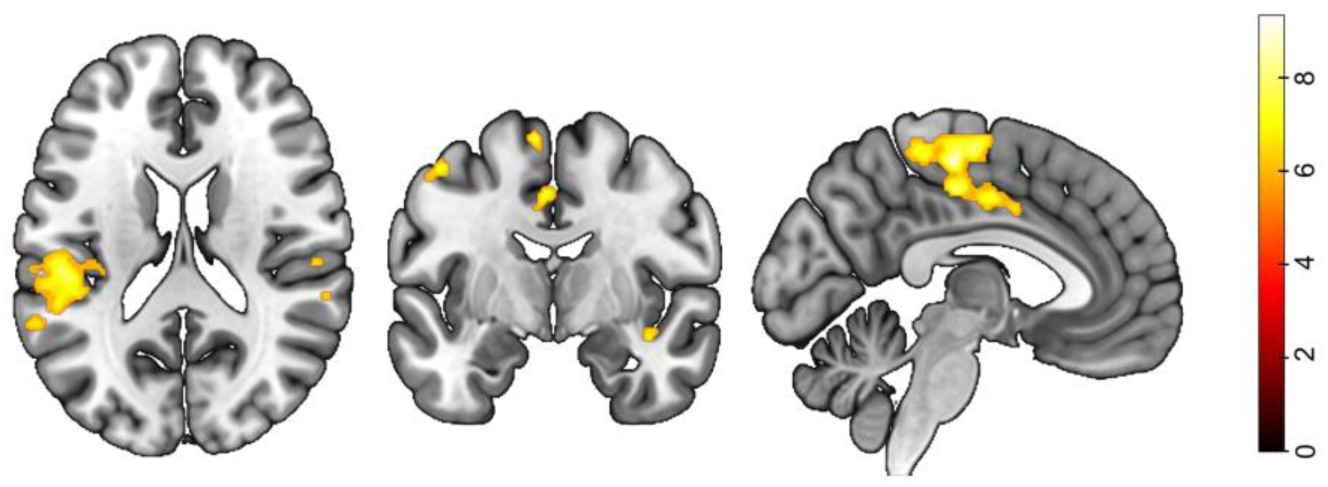
Whole-brain voxel-level family-wise error corrected group-level results (*p < 0.05*) for the contrast Shock > No-Shock during anticipation of choice.

#### 3.5.5. Neural correlates of episodic memory retrieval

The contrast of Remembered > Forgotten across both No-Shock and Shock conditions revealed distributed brain regions with stronger activation for recognized items compared to forgotten items during food stimulus presentation. These regions included the precentral gyrus, postcentral gyrus, calcarine, lingual gyrus, posterior cingulate, and parahippocampal gyrus etc. (vFWE, *p < 0.05*; Figure 9A, Table S7). We observed a significant positive association between the contrast estimates of Remembered > Forgotten and memory accuracy in the OFC (*r*[42] = 0.335, *p* = 0.026; Figure 9B). Time course extracted from the OFC revealed that remembered items induced higher OFC activity than forgotten items (Figure 9C). For the effects of stress on episodic memory retrieval, see Figure S2, Table S8, and Table S9.

**Figure 9.**
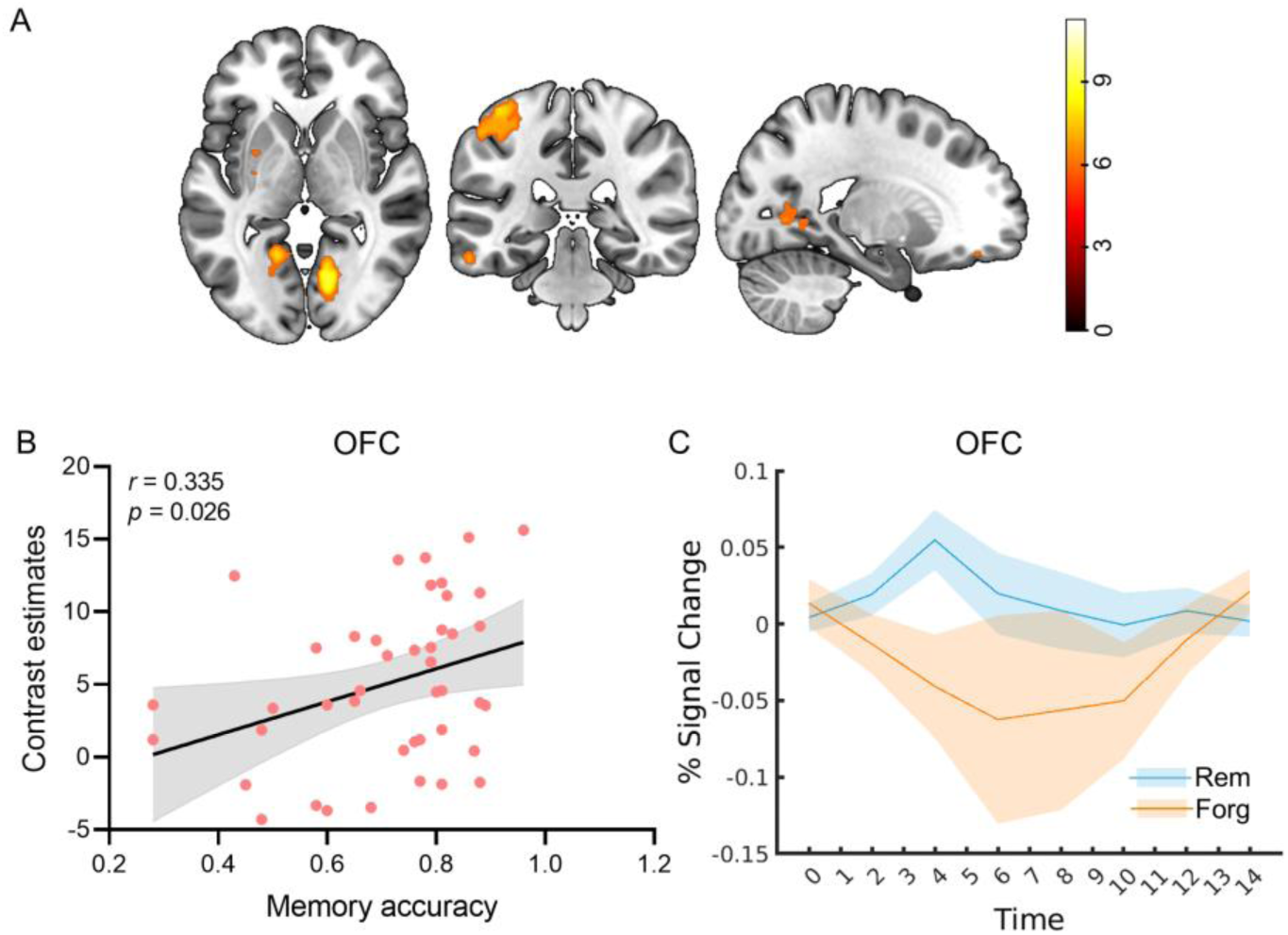
A) Whole-brain voxel-level family-wise error corrected group-level results (*p < 0.05*) for the contrast Remembered > Forgotten; B) A significant positive correlation between the OFC activity during the memory retrieval of food stimulus and memory accuracy; C) Time course of the OFC in the Remembered and Forgotten conditions. Rem, remembered; Forg, forgotten.

## 4. Discussion

This study combined computational modeling, neuroimaging, and behavioral assessments to elucidate the mechanisms by which stress and memory affect choice consistency. Consistent with prior research (Alós-Ferrer & Garagnani, 2021; Krajbich et al., 2010; Polanía et al., 2019), we found that smaller value differences predict lower choice consistency and longer response times, demonstrating the reliability of our experimental paradigm. Additionally, remembered items exhibited higher choice consistency compared to forgotten items. Stress was found to impair both choice consistency and memory retrieval, with stress-induced decreases in memory accuracy positively correlating with decreases in choice reaction times. Using computational modeling, we observed that the drift rate was higher and the decision threshold lower for remembered food items compared to forgotten ones. Additionally, the decision threshold was reduced in the Shock condition compared to the No-Shock condition. Crucially, our study provides evidence for the neural mechanisms influencing choice consistency under varying stress and memory conditions. Our findings demonstrated a positive association between DLPFC activation during the pre-choice anticipation period and choice consistency. Furthermore, we observed a positive association between OFC activation during the memory retrieval of food stimuli and memory accuracy.

### 4.1 Choice consistency and it’s neural mechanisms

Choice consistency is a fundamental aspect of rational decision-making, reflecting the stability and reliability of an individual’s preferences. Our findings indicate that smaller value differences between options correspond to higher levels of choice difficulty. Difficult choices lead to greater inconsistency, with inconsistent choices taking longer to make than consistent ones. This suggests that when participants made choices that deviated from their previously stated preferences, they required more time to decide. One explanation is that this increased reaction time may reflect the cognitive effort involved in reconciling conflicting information or uncertainty. Our fMRI results support this explanation, as the contrast of Inconsistent > Consistent during the choice period revealed activation in regions involved in cognitive control and conflict monitoring, such as the anterior cingulate cortex (Botvinick et al., 2001), DLPFC, superior frontal gyrus, and inferior frontal gyrus (Mansouri et al., 2009). The theory of cognitive dissonance posits that individuals experience cognitive dissonance, characterized by negative emotional arousal due to cognitive conflicts, particularly when making difficult choices between equally attractive items (Festinger, 1957). Kitayama et al. (2013) found that choices between CDs that were close in attractiveness (difficult choices) as opposed to those that were distant (easy choices) resulted in the activation of the dorsal anterior cingulate cortex (dACC), a brain region associated with cognitive conflict (Bush et al., 2000). Another explanation is that participants may have begun preparing during the anticipation period before the choice task. Consequently, trials without prior preparation required more time for decision-making and were more likely to result in inconsistent choices. The fMRI results also support this explanation, with the contrast of Consistent > Inconsistent during the anticipatory choice period revealing activation in the DLPFC. Moreover, anticipatory DLPFC activation is positively correlated with choice consistency. Further research could explore the underlying mechanisms driving these differences in reaction times and investigate various contributing factors.

### 4.2. Influence of memory retrieval on choice consistency

To date, only one study has directly examined the influence of memory processes on choice consistency (Nitsch & Kalenscher, 2021). However, the presence of a floor effect in their data compromised its quality, making it challenging to conclusively evaluate their hypotheses. Our study, to the best of our knowledge, is the second to investigate this topic, aiming to provide more robust and reliable insights. We integrated a recognition memory test with a value-based choice task to examine the impact of memory on choice consistency. Our findings indicate that items remembered by participants demonstrated significantly higher choice consistency and shorter decision reaction time compared to items that were forgotten. Memory has been documented to play an essential role in value-based decision-making (M. N. Shadlen & D. Shohamy, 2016). Many such decisions involve sampling value-relevant evidence from memory to guide the decision-making process. The speed and accuracy of these decisions are often explained by the accumulation of evidence until it reaches a certain threshold. Decision-makers tend to recall their preferences for remembered options both more quickly and accurately. Computational modeling further dissected this effect, revealing that remembered items exhibited a higher drift rate and a lower decision threshold compared to forgotten items. A lower decision threshold represents a smaller distance between two selection boundaries, meaning less evidence is required before making a decision. This can lead to faster decisions. Conversely, a higher drift rate signifies a faster rate of information acquisition, enabling the decision-maker to accumulate evidence more quickly, which often results in faster and more accurate decisions (Ratcliff & McKoon, 2008).

We investigated the neural mechanisms involved in memory retrieval and the impact of stress on this process. Our findings provide evidence for the recruitment of the OFC during memory retrieval. Specifically, we found a significant positive correlation between OFC activity during the retrieval of food-related stimuli and memory accuracy. Furthermore, the temporal dynamics of OFC activation revealed greater activity for remembered items compared to forgotten ones. The OFC plays a significant role in food memory, particularly in encoding and retrieving the sensory and reward-related aspects of food (Brand & Markowitsch, 2006). The OFC processes the sight, smell, taste, and texture of food, integrating these sensory inputs with reward values, which is essential for food intake decisions. Additionally, the OFC’s role in evaluating the healthiness and tastiness of food cues further underscores its importance in food-related decision-making (Londerée & Wagner, 2020). Its anatomical connections to mnemonic regions enable it to interact with other structures involved in memory processes. Research has demonstrated that the OFC is activated when recalling food-related memories, especially those associated with positive experiences or rewards (Morris & Dolan, 2001; Rolls et al., 2021). This activation is linked to the OFC’s connections with other brain regions involved in memory and emotion, such as the amygdala and hippocampus.

### 4.3. Stress impaired both choice consistency and memory retrieval

We observed that acute stress led to a decrease in choice consistency. Additionally, decision response times were shorter, indicating that participants made faster but less rational choices. Computational modeling supports this finding, showing that the decision threshold was lower in the Shock condition compared to the No-Shock condition. Stress has been shown to elicit a switch from an analytic reasoning system to intuitive processes (Yu, 2016).Unlike deliberative thinking, intuitive thinking increases cognitive biases and the likelihood of errors. According to rational choice theory, rational decision-making requires thorough value assessment and comparison of options. When choosing between options of similar value, these deliberative decisions take more time. From this perspective, stress impairs decision quality, negatively affecting value-based choice consistency.

Our results are inconsistent with previous findings, although research on the impact of stress on choice consistency is limited. Cettolin et al. (2020) examined the effect of physiological stress on choice consistency in economic decisions and found that economic rationality, defined as consistency with the Generalized Axiom of Revealed Preference (GARP), is not impaired by physiological stress. Nitsch et al. (2021) investigated the impact of acute stress on choice consistency in a time dependent fashion using a food choice task. Their findings provided strong evidence against the influence of acute stress on choice consistency. However, they found exploratory evidence suggesting that higher exposure to chronic stress is associated with decreased choice consistency. Neither study observed an effect of acute stress on choice consistency. One possible reason for this is that participants had ample decision-making time in the experiments, allowing for thorough valuation and decision. Without time pressure, rational decisions are more likely. However, decisions often need to be made within seconds. According to the available literature, our study is the first to examine the impact of acute stress on choice consistency under time pressure. Furthermore, rapid decision-making under these conditions is well-suited for investigating cognitive mechanisms using the Drift Diffusion Model (DDM).

Acute stress during decision-making significantly influences brain activation patterns, highlighting its complex impact on neural dynamics. Specifically, during the choice period, stress reduced activation in the postcentral gyrus, precentral gyrus, fusiform gyrus, and parahippocampal gyrus, which are typically associated with sensorimotor processing and memory. In contrast, stress enhanced activation in regions including the precuneus, superior temporal gyrus, supramarginal gyrus, insula, operculum, and lingual gyrus, which are often implicated in emotional salience. These findings suggest that acute stress may impair memory and motor functions while enhancing emotional and sensory integration during decision-making. During the anticipation period preceding the choice, stress enhanced activation in the medial frontal gyrus, postcentral gyrus, middle cingulate, superior temporal gyrus, parietal operculum, supramarginal gyrus, precentral gyrus, superior parietal lobule, insula, rolandic operculum, and middle temporal gyrus. These regions are involved in attentional control, sensory processing, emotional evaluation, and motor preparation. The increased activity suggests that acute stress primes the brain for heightened sensory and cognitive readiness in anticipation of a decision, possibly as an adaptive response to environmental demands. The distinct neural activation patterns during the anticipation and choice periods highlight the multifaceted influence of acute stress on neural dynamics. Stress appears to reallocate cognitive and neural resources, dampening activity in regions essential for detailed spatial and sensorimotor processing while enhancing activation in areas involved in emotional and attentional prioritization. Regions such as the precuneus and insula, which showed increased activation, are known to be associated with heightened interoceptive awareness and affective processing under stress. Meanwhile, the reduced activation in memory-related areas like the parahippocampal gyrus may explain the observed memory impairments under stress.

In line with numerous prior studies (Gagnon & Wagner, 2016; Gagnon et al., 2019; Wolf, 2017), our findings demonstrate that acute stress impairs memory retrieval, as evidenced by a significant decrease in recognition accuracy. The effects of acute stress on memory retrieval revealed activation changes across various brain regions. Specifically, stress decreased activation in the insula, inferior frontal gyrus, thalamus, claustrum, precuneus, caudate, lentiform nucleus, medial frontal gyrus, cingulate gyrus, supramarginal gyrus, inferior parietal lobule, postcentral gyrus, culmen, and anterior cingulate. These areas are involved in various cognitive functions, including attention, sensory processing, executive control, and memory retrieval. Conversely, stress enhanced activation in the amygdala, middle frontal gyrus, hippocampus, entorhinal area, parahippocampal gyrus, superior frontal gyrus, and superior temporal gyrus. These regions are associated with emotional processing, memory encoding, and retrieval, suggesting that stress heightens the brain’s response to emotionally salient and memory-related tasks. These findings underscore the complex and widespread impact of acute stress on neural mechanisms underlying memory retrieval. The enhanced activation in regions related to emotional and memory processing may reflect an adaptive response to stress, prioritizing the encoding and retrieval of emotionally relevant information (Schwabe et al., 2022). Stress-induced activation of the amygdala may skew memory retrieval towards emotional or irrelevant details, thereby hindering access to neutral or precise episodic memories. However, the concurrent decrease in activation in regions associated with broader cognitive functions indicates that this prioritization comes at the cost of reduced efficiency in other cognitive domains.

Overall, stress impairs both memory and choice consistency. Additionally, we observed a positive correlation between stress-induced decreases in memory accuracy and reductions in choice reaction times. This suggests that stress may impair choice consistency by disrupting memory retrieval processes. However, further research is needed to validate this conclusion.

## 5. Conclusions

This study combined computational modeling, neuroimaging, and behavioral assessments to elucidate the mechanisms by which stress and memory affect choice consistency. Our findings reveal that accurate memory retrieval leads to more consistent choices. Acute stress was found to impair both choice consistency and memory retrieval, suggesting that stress may disrupt choice consistency by impairing the retrieval of option values from memory. Importantly, our results highlight the neural mechanisms underlying choice consistency under varying levels of stress and memory conditions. These findings provide novel insights into the role of stress and memory in decision-making, offering a more nuanced understanding of the neural and cognitive processes that govern choice behavior.

## Conflict of Interest

The authors declare no conflict of interest.

## Acknowledgments

This work was supported by the National Natural Science Foundation of China (32200909); the Shenzhen-Hong Kong Institute of Brain Science-Shenzhen Fundamental Research Institutions (2024SHIBS0003); the University stability support program of Shenzhen (Project 20220810160130001).

## Author contributions

Fei Xin: conceived and designed the study, analyzed the data, wrote the paper. Jialuo Lai: analyzed the data, wrote the paper. Manru Guo: collected the data. Qingfei Chen: wrote the paper. Jianhui Wu: wrote the paper.

## Notes

### Competing Interest Statement

The authors have declared no competing interest.

