## Supplementary Materials for "The Impact of Memory and Stress on Choice Consistency"

### Affiliations

### *Corresponding author:


### The file includes:

Tables S1-S9; Figure S1-S2.

**Table S1**. Results of model comparisons in hierarchical drift diffusion model analyses.

| Model No. | Model | DIC | WAIC | LOOCV |
|---|---|---|---|---|
| 1 | *v* ~ Memory<br>α ~ Memory + Stress | 10497.65(1) | 10526.82(1) | 10528.34(1) |
| 2 | *v* ~ Stress<br>α ~ Memory * Stress | 10501.34(2) | 10557.82(6) | 10559.83(6) |
| 3 | *v* ~ Memory + Stress<br>α ~ Memory + Stress | 10518.44(3) | 10544.06(3) | 10545.95(3) |
| 4 | *v* ~ Stress<br>α ~ Memory + Stress | 10526.56(4) | 10567.94(8) | 10570.10(8) |
| 5 | *v* ~ Stress<br>α ~ Memory * Stress | 10539.33(5) | 10579.40(9) | 10581.23(9) |
| 6 | *v* ~ Memory<br>α ~ Memory * Stress | 10543.25(6) | 10581.89(10) | 10583.99(10) |
| 7 | *v* ~ Memory + Stress<br>α ~ Stress | 10549.98(7) | 10552.45(5) | 10553.67(5) |
| 8 | *v* ~ Stress<br>α ~ Stress | 10555.25(8) | 10567.74(7) | 10569.10(7) |
| 9 | *v* ~ Memory * Stress<br>α ~ Stress * Memory | 10555.64(9) | 10551.12(4) | 10553.07(4) |
| 10 | *v* ~ Memory: Stress<br>α ~ Memory: Stress | 10578.66(10) | 10543.00(2) | 10545.59(2) |
| 11 | *v* ~ Memory + Stress<br>α ~ Memory | 10657.08(11) | 10679.28(11) | 10680.63(11) |
| 12 | *v* ~ Memory * Stress<br>α ~ Memory | 10661.08(12) | 10714.57(13) | 10716.32(13) |
| 13 | *v* ~ Memory<br>α ~ Memory | 10676.11(13) | 10690.57(12) | 10690.49(12) |
| 14 | *v* ~ Memory: Stress | 10902.33(14) | 10937.91(14) | 10939.43(14) |
| 15 | α ~ Memory: Stress | 12189.84(15) | 12148.41(15) | 12151.40(15) |

DIC, WAIC, and LOOCV are expressed as values (rank). Lower values of DIC, WAIC, and LOOCV indicate a better model fit. Since DIC favors models with greater complexity, we used WAIC and LOOCV as additional evaluation metrics to measure model robustness and guard against overfitting. Therefore, we selected Model 1 as the best-fitting model. Detailed expressions related to model modeling can be found on the Pasty official website. Each model uses the Markov Chain Monte Carlo method to estimate the joint posterior distribution for all parameters with 5000 samples, discarding the first 1000 burn-in samples to allow for convergence. DIC, deviance information criterion; WAIC, widely applicable information criterion; LOOCV, Pareto-smoothed importance sampling leave-one-out cross-validation; *v*, drift rate; α, threshold.

**Table S2.** Activation Table for GLM Analysis of choice (Inconsistent > Consistent).

| Region | Side | Cluster size | Peak Z | MNI Coordinates | | |
|---|---|---|---|---|---|---|
| | | | | x | y | z |
| Anterior Cingulate/Medial Frontal Gyrus/Superior Frontal Gyrus | L | 679 | 4.95 | -10 | 30 | 22 |
| Supplementary Motor Area | L | | 4.70 | 0 | 22 | 38 |
| Supplementary Motor Area | L | | 4.26 | -8 | 10 | 46 |
| Angular/Inferior Parietal Lobule/Precuneus | L | 521 | 4.93 | -36 | -56 | 36 |
| Superior Parietal Lobule | L | | 3.97 | -34 | -42 | 44 |
| Superior Parietal Lobule | L | | 3.66 | -26 | -62 | 44 |
| Inferior Frontal Gyrus/Middle Frontal Gyrus | L | 676 | 4.54 | -52 | 14 | 30 |
| Middle Frontal Gyrus | L | | 4.16 | -44 | 32 | 34 |
| Middle Frontal Gyrus | L | | 4.10 | -40 | 6 | 38 |
| Cerebellum | R | 519 | 4.46 | 32 | -60 | -32 |
| Cerebellum | R | | 4.12 | 40 | -58 | -32 |
| Cerebellum | R | | 4.10 | 24 | -68 | -26 |
| Superior Frontal Gyrus/Middle Frontal Gyrus | L | 244 | 4.33 | -28 | 4 | 66 |
| Superior Frontal Gyrus | L | | 4.18 | -24 | 16 | 66 |
| Middle Frontal Gyrus | L | 220 | 4.14 | -46 | 54 | 8 |
| Middle Frontal Gyrus | L | | 4.13 | -40 | 48 | 22 |
| Middle Frontal Gyrus | L | | 3.49 | -34 | 58 | 18 |
| Middle Frontal Gyrus/Inferior Frontal Gyrus | R | 166 | 4.11 | 50 | 18 | 44 |
| Middle Frontal Gyrus | R | | 3.83 | 58 | 14 | 36 |
| Superior Parietal Lobule/Precuneus | R | 165 | 4.03 | 26 | -74 | 52 |
| Superior Parietal Lobule/Precuneus | R | | 3.54 | 10 | -68 | 56 |
| Superior Parietal Lobule/Precuneus | R | | 3.46 | 12 | -76 | 58 |
| Cerebellum | L | 244 | 3.99 | -34 | -60 | -30 |
| Cerebellum | L | | 3.73 | -38 | -52 | -32 |
| Cerebellum | L | | 3.73 | -46 | -58 | -36 |

GLM, general linear model; L, left; MNI, Montreal Neurological Institute; R, right. $p < 0.05$, cluster-level family-wise error (FWE) corrected.

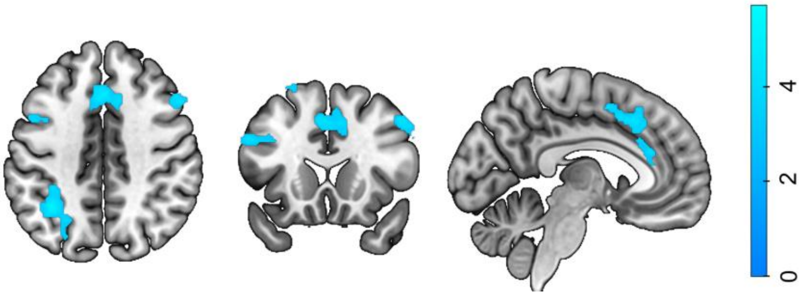


**Figure S1**. Whole-brain cluster-level family-wise error corrected group-level results ($p < 0.05$) for the contrast Inconsistent > Consistent during choice period.

**Table S3**. Activation Table for GLM Analysis of stress effects on choice (Shock > No-Shock).

| **Region** | **Side** | **Cluster size** | **Peak Z** | **MNI Coordinates** | | |
|---|---|---|---|---|---|---|
| | | | | **x** | **y** | **z** |
| Precuneus/Precentral gyrus/Superior Frontal Gyrus | L | 1391 | 5.08 | -8 | -22 | 80 |
| Postcentral gyrus | L | | 4.61 | -12 | -40 | 78 |
| Precuneus | R | | 4.46 | 2 | -50 | 52 |
| SupraMarginal gyrus/Superior Temporal Gyrus/Inferior Parietal Lobule | R | 854 | 4.70 | 62 | -38 | 26 |
| Superior Temporal Gyrus | R | | 4.50 | 68 | -18 | 14 |
| SupraMarginal gyrus | R | | 4.44 | 70 | -30 | 24 |
| Insula | L | 206 | 4.40 | -22 | -26 | 16 |
| Operculum/Insula | L | | 4.39 | -34 | -18 | 24 |
| Operculum/Insula | L | | 3.89 | -32 | -8 | 24 |
| Superior Temporal Gyrus/SupraMarginal gyrus | L | 439 | 4.22 | -64 | -36 | 20 |
| SupraMarginal gyrus | L | | 4.06 | -66 | -44 | 28 |
| SupraMarginal gyrus | L | | 3.90 | -56 | -40 | 32 |
| Lingual Gyrus/Cuneus | L | 444 | 4.14 | 0 | -86 | -26 |
| Lingual Gyrus | L | | 3.89 | -4 | -72 | -4 |
| Lingual Gyrus | R | | 3.67 | 4 | -72 | -6 |
| Precuneus | R | 385 | 4.14 | 10 | -68 | 34 |
| Precuneus | L | | 3.72 | -8 | -80 | 38 |
| Cuneus | R | | 3.51 | 4 | -84 | 38 |

GLM, general linear model; L, left; MNI, Montreal Neurological Institute; R, right. $p < 0.05$, cluster-level family-wise error (FWE) corrected.

**Table S4**. Activation Table for GLM Analysis of stress effects on choice (No-Shock >Shock).

| **Region** | **Side** | **Cluster size** | **Peak Z** | **MNI Coordinates** | | |
|---|---|---|---|---|---|---|
| | | | | **x** | **y** | **z** |
| Postcentral gyrus/Precentral Gyrus/Inferior Parietal Lobule | L | 1085 | 5.42 | -52 | -22 | 56 |
| Postcentral gyrus | L | | 4.95 | -56 | -14 | 48 |
| Postcentral gyrus | L | | 4.68 | -46 | -18 | 52 |
| Precentral Gyrus/Postcentral Gyrus | R | 646 | 4.79 | 44 | -26 | 68 |
| Postcentral gyrus | R | | 4.75 | 52 | -22 | 62 |
| Postcentral gyrus | R | | 4.23 | 38 | -32 | 50 |
| Fusiform Gyrus | L | 229 | 4.60 | -34 | -66 | -16 |
| Fusiform Gyrus | L | | 4.15 | -36 | -52 | -18 |
| Fusiform Gyrus | L | | 3.53 | -28 | -44 | -18 |
| Fusiform Gyrus/Parahippocampa Gyrus | L | 229 | 4.47 | -34 | -30 | -22 |
| Parahippocampa Gyrus | L | | 4.30 | -30 | -22 | -30 |
| Fusiform Gyrus | L | | 4.08 | -38 | -24 | -26 |
| Fusiform Gyrus/Parahippocampa Gyrus/Hippocampus | R | 377 | 4.45 | 30 | -18 | -22 |
| Fusiform Gyrus | R | | 4.41 | 30 | -40 | -22 |
| Fusiform Gyrus | R | | 3.99 | 34 | -48 | -24 |

GLM, general linear model; L, left; MNI, Montreal Neurological Institute; R, right. $p < 0.05$, cluster-level family-wise error (FWE) corrected.

**Table S5**. Activation Table for GLM Analysis of anticipatory response before choice (Consistent > Inconsistent).

| **Region** | **Side** | **Cluster size** | **Peak Z** | **MNI Coordinates** | | |
|---|---|---|---|---|---|---|
| | | | | **x** | **y** | **z** |
| Middle Frontal Gyrus/Superior Frontal Gyrus | L | 258 | 4.78 | -38 | 36 | 28 |
| Superior Frontal Gyrus | L | | 3.94 | -22 | 52 | 26 |
| SupraMarginal gyrus/Inferior Parietal Lobule/Insula | R | 124 | 4.48 | 58 | -36 | 24 |
| Middle Cingulate | L | 249 | 4.30 | -10 | 10 | 32 |
| Middle Cingulate | R | | 4.02 | 10 | -2 | 34 |
| Middle Cingulate | R | | 3.77 | 4 | 2 | 42 |
| Cerebellum | R | 174 | 4.24 | 48 | -72 | -40 |
| Cerebellum | R | | 3.57 | 44 | -54 | -42 |
| SupraMarginal gyrus/Inferior Parietal Lobule | L | 188 | 4.05 | -58 | -42 | 28 |
| SupraMarginal gyrus/Inferior Parietal Lobule | L | | 3.75 | -58 | -32 | 26 |
| Cerebellum | L | 122 | 3.93 | -36 | -78 | -32 |
| Cerebellum | L | | 3.69 | -42 | -78 | -40 |

GLM, general linear model; L, left; MNI, Montreal Neurological Institute; R, right. $p < 0.05$, cluster-level family-wise error (FWE) corrected.

**Table S6**. Activation Table for GLM Analysis of stress effects on anticipatory response before choice (Shock > No-Shock).

| **Region** | **Side** | **Cluster size** | **Peak Z** | **MNI Coordinates** | | |
|---|---|---|---|---|---|---|
| | | | | **x** | **y** | **z** |
| Paracentral Lobule/Medial Frontal Gyrus/Middle Cingulate/Precuneus | L | 1076 | 6.91 | -2 | -22 | 58 |
| Postcentral Gyrus | L | | 6.80 | -12 | -34 | 66 |
| Middle Cingulate | R | | 6.47 | 10 | -12 | 42 |
| Superior Temporal Gyrus/Insula/Rolandic_Oper/Inferior Parietal Lobule | L | 477 | 6.29 | -48 | -28 | 18 |
| Parietal operculum | L | | 6.09 | -52 | -40 | 24 |
| SupraMarginal Gyrus | L | | 5.83 | -60 | -48 | 20 |
| Precentral Gyrus/Postcentral Gyrus | R | 94 | 6.08 | 20 | -32 | 64 |
| Postcentral Gyrus | R | | 5.55 | 28 | -30 | 64 |
| Superior Parietal Lobule | R | | 5.45 | 22 | -40 | 62 |
| Insula | L | 31 | 6.02 | -36 | 2 | 10 |
| Precentral Gyrus | L | 57 | 5.98 | -46 | -6 | 54 |
| Precentral Gyrus | L | | 5.24 | -42 | -8 | 62 |
| Rolandic operculum | R | 18 | 5.70 | 52 | 4 | 4 |
| Insula | R | 15 | 5.68 | 40 | -6 | -14 |
| Rolandic operculum | L | 35 | 5.62 | -60 | 2 | 8 |
| Rolandic operculum | R | 14 | 5.62 | 56 | -24 | 20 |
| Middle Cingulate | R | 16 | 5.54 | 10 | -2 | 40 |
| Superior Temporal Gyrus | R | 9 | 5.35 | 60 | -36 | 18 |
| Insula | L | 1 | 5.32 | -36 | 8 | -2 |
| Superior Parietal Lobule | R | 6 | 5.28 | 16 | -48 | 66 |

| | | | | | | |
|---|---|---|---|---|---|---|
| Insula | R | 2 | 5.27 | 34 | -18 | 16 |
| Cerebellum | R | 2 | 5.27 | 2 | -66 | -40 |
| Middle Temporal Gyrus | L | 7 | 5.27 | -54 | -56 | 4 |
| SupraMarginal Gyrus/Angular gyrus | R | 2 | 5.21 | 60 | -44 | 28 |
| Middle Temporal Gyrus | L | 3 | 5.15 | -44 | -56 | 8 |
| Paracentral Lobule | R | 1 | 5.14 | 6 | -34 | 54 |
| Cerebrum | R | 1 | 5.11 | 36 | -18 | -4 |
| Superior Temporal Gyrus | R | 1 | 5.10 | 62 | -36 | 10 |
| SupraMarginal | R | 1 | 5.08 | 46 | -30 | 24 |
| Postcentral Gyrus | L | 1 | 5.05 | -24 | -42 | 60 |

GLM, general linear model; L, left; MNI, Montreal Neurological Institute; R, right. $p < 0.05$, voxel-level family-wise error (FWE) corrected.

**Table S7**. Activation Table for GLM Analysis of memory retrieval (Remembered > Forgotten).

| **Region** | **Side** | **Cluster size** | **Peak Z** | **MNI Coordinates** | | |
|---|---|---|---|---|---|---|
| | | | | **x** | **y** | **z** |
| Precentral Gyrus/Postcentral Gyrus/Inferior Parietal Lobule | L | 1605 | 7.72 | -32 | -22 | 48 |
| Precentral Gyrus/Postcentral Gyrus | L | | 7.45 | -34 | -22 | 58 |
| Postcentral Gyrus | L | | 6.74 | -50 | -18 | 56 |
| Calcarine/Lingual Gyrus/Posterior Cingulate | R | 701 | 7.23 | 14 | -76 | 4 |
| Lingual Gyrus | R | | 7.17 | 10 | -66 | -6 |
| Cerebellum exterior | R | | 6.24 | 16 | -58 | -22 |
| Calcarine/Posterior Cingulate/Precuneus/Lingual Gyrus/Parahippocampal gyrus | L | 450 | 6.47 | -14 | -60 | 8 |
| Lingual Gyrus/Precuneus | L | | 6.23 | -16 | -56 | 0 |
| Calcarine | L | | 5.70 | -20 | -68 | 4 |
| Cuneus/Precuneus | L | 58 | 5.89 | -14 | -74 | 34 |
| Precuneus | L | | 5.58 | -4 | -78 | 32 |
| Supplementary Motor Area/ Middle Cingulate/Medial Frontal Gyrus | L | 35 | 5.86 | -4 | -16 | 50 |
| Rolandic Operculum/Insula | L | 18 | 5.79 | -44 | -22 | 18 |
| Inferior Temporal Gyrus/Middle Temporal Gyrus | L | 33 | 5.78 | -58 | -34 | -20 |
| Middle Frontal Gyrus | R | 17 | 5.74 | 46 | 52 | 16 |
| Precuneus | L | 20 | 5.65 | -10 | -68 | 54 |
| Inferior Parietal Lobule | L | 21 | 5.65 | -38 | -56 | 50 |
| Cuneus | R | 39 | 5.64 | 18 | -84 | 24 |
| Cuneus | R | | 5.06 | 12 | -82 | 30 |
| Inferior Temporal Gyrus/Middle Temporal Gyrus | L | 13 | 5.47 | -54 | -54 | -14 |
| Putamen | L | 10 | 5.41 | -30 | 0 | -2 |
| Middle Temporal Gyrus/Inferior Temporal Gyrus | L | 7 | 5.34 | -60 | -22 | -20 |
| Anterior orbital gyrus/ Medial orbital gyrus | L | 5 | 5.30 | -20 | 44 | -16 |
| Middle Frontal Gyrus | L | 8 | 5.30 | -40 | 40 | 30 |
| Cerebellum_Crus1 | R | 3 | 5.26 | 42 | -70 | -36 |
| Inferior Parietal Lobule | L | 3 | 5.25 | -54 | -38 | 40 |
| Lingual Gyrus /Parahippocampal gyrus | R | 2 | 5.20 | 22 | -56 | -10 |
| Putamen | L | 3 | 5.20 | -30 | -10 | 0 |
| Calcarine | L | 2 | 5.19 | 0 | -70 | 20 |

| | | | | | | |
|---|---|---|---|---|---|---|
| Inferior Temporal Gyrus | R | 2 | 5.15 | 46 | -54 | -10 |
| Cerebellum_8 | R | 1 | 5.13 | 14 | -70 | -42 |
| Inferior Parietal Lobule | L | 1 | 5.08 | -40 | -30 | 42 |

GLM, general linear model; L, left; MNI, Montreal Neurological Institute; R, right. $p < 0.05$, voxel-level family-wise error (FWE) corrected.

**Table S8**. Activation Table for GLM Analysis of stress effects on memory retrieval (Shock > No-Shock).

| **Region** | **Side** | **Cluster size** | **Peak Z** | **MNI Coordinates** | | |
|---|---|---|---|---|---|---|
| | | | | **x** | **y** | **z** |
| Middle Frontal Gyrus | L | 165 | 5.04 | -24 | 36 | -14 |
| Middle Frontal Gyrus | L | | 3.58 | -38 | 38 | -18 |
| Middle Frontal Gyrus | L | | 3.34 | -36 | 44 | -12 |
| Amygdala | R | 223 | 4.99 | 28 | -4 | -26 |
| Hippocampus/Amygdala | R | | 4.39 | 22 | -12 | -20 |
| Entorhinal area/Amygdala | R | | 4.29 | 22 | 0 | -22 |
| Hippocampus/Parahippocampal Gyrus /Amygdala | L | 126 | 4.68 | -22 | -16 | -20 |
| Hippocampus/Amygdala | L | | 3.38 | -16 | -8 | -24 |
| Parahippocampal Gyrus | L | | 3.21 | -22 | -24 | -22 |
| Superior Frontal Gyrus | R | 134 | 4.55 | 4 | 72 | 16 |
| Superior Frontal Gyrus | R | | 3.46 | 0 | 66 | 24 |
| Superior Temporal Gyrus | R | 120 | 4.39 | 60 | -4 | -2 |
| Superior Temporal Gyrus | R | | 3.46 | 58 | -12 | 4 |

GLM, general linear model; L, left; MNI, Montreal Neurological Institute; R, right. $p < 0.05$, cluster-level family-wise error (FWE) corrected.

**Table S9**. Activation Table for GLM Analysis of stress effects on memory retrieval (No-Shock > Shock).

| **Region** | **Side** | **Cluster size** | **Peak Z** | **MNI Coordinates** | | |
|---|---|---|---|---|---|---|
| | | | | **x** | **y** | **z** |
| Insula | R | 1436 | 6.71 | 30 | 22 | -6 |
| Inferior Frontal Gyrus | R | | 5.60 | 48 | 10 | 14 |
| Inferior Frontal Gyrus | R | | 5.18 | 52 | 22 | 6 |
| Thalamus | L | 803 | 6.17 | -6 | -28 | -6 |
| Thalamus | L | | 5.43 | -6 | -18 | 8 |
| Thalamus | R | | 4.80 | 6 | -20 | 10 |
| Superior Frontal Gyrus | R | 295 | 5.41 | 6 | 12 | 54 |
| Superior Frontal Gyrus | L | | 4.65 | -6 | 20 | 54 |
| Superior Frontal Gyrus | L | | 3.29 | -8 | 14 | 60 |
| Inferior Frontal Gyrus | L | 1050 | 5.35 | -28 | 16 | -14 |
| Inferior Frontal Gyrus | L | | 5.11 | -50 | 18 | 4 |
| Claustrum | L | | 4.86 | -28 | 10 | 8 |
| Cuneus | L | 164 | 5.20 | -10 | -68 | 30 |
| Cuneus | L | | 3.59 | -4 | -78 | 38 |
| Precuneus | L | | 3.37 | -18 | -60 | 24 |
| Precuneus | R | 287 | 5.20 | 14 | -60 | 32 |
| Cuneus | R | | 5.06 | 8 | -70 | 34 |

| | | | | | | |
|---|---|---|---|---|---|---|
| Caudate | L | 125 | 4.80 | -10 | 6 | 2 |
| Lentiform Nucleus | L | | 3.82 | -14 | 10 | -4 |
| Medial Frontal Gyrus | R | 224 | 4.65 | 4 | 34 | 34 |
| Cingulate Gyrus | R | | 4.27 | 10 | 26 | 32 |
| Cingulate Gyrus | L | | 4.09 | -8 | 18 | 30 |
| Supramarginal Gyrus | L | 252 | 4.65 | -52 | -38 | 28 |
| Inferior Parietal Lobule | L | | 4.14 | -66 | -36 | 22 |
| Postcentral Gyrus | L | | 3.14 | -58 | -28 | 22 |
| Caudate | R | 152 | 4.63 | 10 | 8 | 4 |
| Caudate | R | | 3.56 | 14 | 14 | 0 |
| Culmen | R | 92 | 4.60 | 0 | -52 | -24 |
| Cerebellar Tonsil | L | | 4.19 | 0 | -56 | -36 |
| Anterior Cingulate | R | 149 | 4.32 | 10 | 38 | 4 |
| Anterior Cingulate | R | | 4.27 | 10 | 34 | 16 |

GLM, general linear model; L, left; MNI, Montreal Neurological Institute; R, right. $p < 0.05$, cluster-level family-wise error (FWE) corrected.

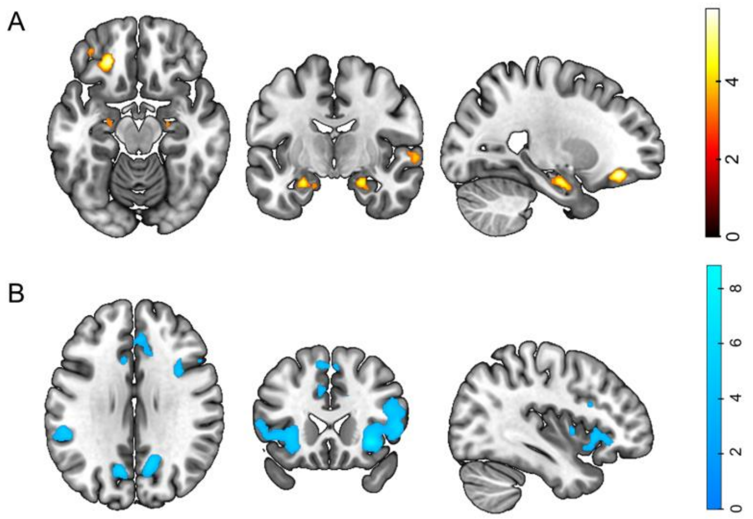


**Figure S2**. The effects of stress on episodic memory retrieval. Whole-brain cluster-level family-wise error corrected group-level results ($p < 0.05$) for the contrast A) Shock > No-Shock and B) No-Shock > Shock during memory retrieval.
